# Structure-Guided Mutagenesis Alters Deubiquitinating Activity and Attenuates Pathogenesis of a Murine Coronavirus

**DOI:** 10.1101/782409

**Authors:** Xufang Deng, Yafang Chen, Anna M. Mielech, Matthew Hackbart, Kristina R. Kesely, Robert C. Mettelman, Amornrat O’Brien, Mackenzie E. Chapman, Andrew D. Mesecar, Susan C. Baker

## Abstract

Coronaviruses express a multifunctional papain-like protease, termed PLP2. PLP2 acts as a protease that cleaves the viral replicase polyprotein, and a deubiquitinating (DUB) enzyme which removes ubiquitin moieties from ubiquitin-conjugated proteins. Previous *in vitro* studies implicated PLP2 DUB activity as a negative regulator of the host interferon (IFN) response, but the role of DUB activity during virus infection was unknown. Here, we used X-ray structure-guided mutagenesis and functional studies to identify amino acid substitutions within the ubiquitin-binding surface of PLP2 that reduced DUB activity without affecting polyprotein processing activity. We engineered a DUB mutation (Asp1772 to Ala) into a murine coronavirus and evaluated the replication and pathogenesis of the DUB mutant virus (DUBmut) in cultured macrophages and in mice. We found that the DUBmut virus replicates similarly as the wild-type virus in cultured cells, but the DUBmut virus activates an IFN response at earlier times compared to the wild-type virus infection in macrophages, consistent with DUB activity negatively regulating the IFN response. We compared the pathogenesis of the DUBmut virus to the wild-type virus and found that the DUBmut-infected mice had a statistically significant reduction (p<0.05) in viral titer in livers and spleens at day 5 post-infection, albeit both wild-type and DUBmut virus infections resulted in similar liver pathology. Overall, this study demonstrates that structure-guided mutagenesis aids the identification of critical determinants of PLP2-ubiquitin complex, and that PLP2 DUB activity plays a role as an interferon antagonist in coronavirus pathogenesis.

**Importance:** Coronaviruses employ a genetic economy by encoding multifunctional proteins that function in viral replication and also modify the host environment to disarm the innate immune response. The coronavirus papain-like protease 2 (PLP2) domain possesses protease activity, which cleaves the viral replicase polyprotein, and also DUB activity (de-conjugating ubiquitin/ubiquitin-like molecules from modified substrates) using identical catalytic residues. To separate the DUB activity from the protease activity, we employed a structure-guided mutagenesis approach and identified residues that are important for ubiquitin-binding. We found that mutating the ubiquitin-binding residues results in a PLP2 that has reduced DUB activity but retains protease activity. We engineered a recombinant murine coronavirus to express the DUB mutant and showed that the DUB mutant virus activated an earlier type I interferon response in macrophages and exhibited reduced pathogenesis in mice. The results of this study demonstrate that PLP2/DUB is an interferon antagonist and a virulence trait of coronaviruses.

## Introduction

Coronaviruses (CoVs) are members of the order *Nidovirales*, which includes enveloped viruses with large (∼30kb), positive-sense single-stranded RNA genomes that yield a characteristic nested set of subgenomic mRNAs during replication in the cytoplasm of infected cells (1, 2). The genome organization for coronaviruses is highly conserved, with the 5’-most two-thirds of the genome encoding the replicase polyprotein, followed by sequences encoding the canonical structural proteins: spike, envelope, membrane, and nucleocapsid. Many CoVs contain accessory genes, which are interspersed among the genes for the structural proteins. Although these accessory genes are not necessarily required for virus replication and are, in general, not highly conserved within the virus family, many encode proteins that regulate the host response (3). Interestingly, coronavirus replicase proteins, which are highly conserved, can also act as antagonists to block or delay the host innate immune response to infection (1, 4–8). That a slew of coronavirus-encoded accessory and non-accessory proteins have been shown to shape the host antiviral response suggests that viral-mediated subversion of host defenses is an important element of infection. Here, we focus on the viral protease/deubiquitinase (DUB) with the goal of assessing the role of DUB activity in shaping the pathogenesis of mouse hepatitis virus (MHV), a model murine coronavirus.

Coronavirus proteases are essential for viral replication as they are responsible for processing the replicase polyprotein (2, 9–11). Murine coronavirus MHV encodes three proteases: two papain-like proteases (PLP1 and PLP2) and one chymotrypsin-like protease (3CLpro, also termed Mpro). MHV PLP2 is similar to the single papain-like protease (termed PLpro) of Severe Acute Respiratory Syndrome Coronavirus (SARS-CoV) and Middle East Respiratory Syndrome Coronavirus (MERS-CoV) (9, 12–16). We and others have revealed that PLP2 or PLpro of multiple coronaviruses, including MHV, are multifunctional, not only capable of cleaving the viral polyprotein but also possessing deubiquitinase (DUB) and deISGylating (de-conjugating ISG15 molecule from modified substrates) activities (9, 12, 13, 15–27). However, it has been challenging to study the effects of a mutated DUB on virus replication and pathogenesis because the protease and DUB activities share the same catalytic site, disruption of which is lethal for virus replication. Therefore, it was necessary for us to identify residue(s) within the MHV protease/DUB-ubiquitin binding surface that can be mutated to result in reduced DUB activity without affecting polyprotein processing. In this report, we describe the X-ray structure-guided identification of such residues and evaluate the replication, IFN antagonistic effect, and pathogenesis of a recombinant DUB mutant MHV.

## Results

### Structure-guided identification of PLP2 residues interacting with ubiquitin

To investigate the role of viral deubiquitinating activity in coronavirus replication and pathogenesis, we first needed to identify amino acid residues that, when mutated, result in reduced DUB activity while preserving the enzyme’s deISGylating and protease activities, the latter being necessary for viral replication. X-ray structural studies of SARS- and MERS-CoV papain-like protease/DUBs co-crystalized with ubiquitin modified at the c-terminus with a covalent warhead (aldehyde, 3-bromopropylamine, or propargylamine) allowed for identification of residues that are important for direct interaction with ubiquitin (17, 18, 20). Here, we took a slightly different approach and mutated the catalytic cysteine (C1716) of MHV PLP2 to a serine residue and co-crystallized it with free ubiquitin. The X-ray structure of MHV PLP2 (C1716S) in complex with ubiquitin was determined to a resolution of 1.85 Å and an overall R_*free*_ = 19.6% and R_*work*_ = 15.8% (PDB: 5WFI). The overall structure of the MHV PLP2-ubiquitin complex is similar to other PLP2/PLpro ubiquitin-bound structures (Fig. 1A). Ubiquitin binds within the palm region and is gripped by the zinc-fingers motif while the C-terminus extends into the active site.

**Figure 1.**
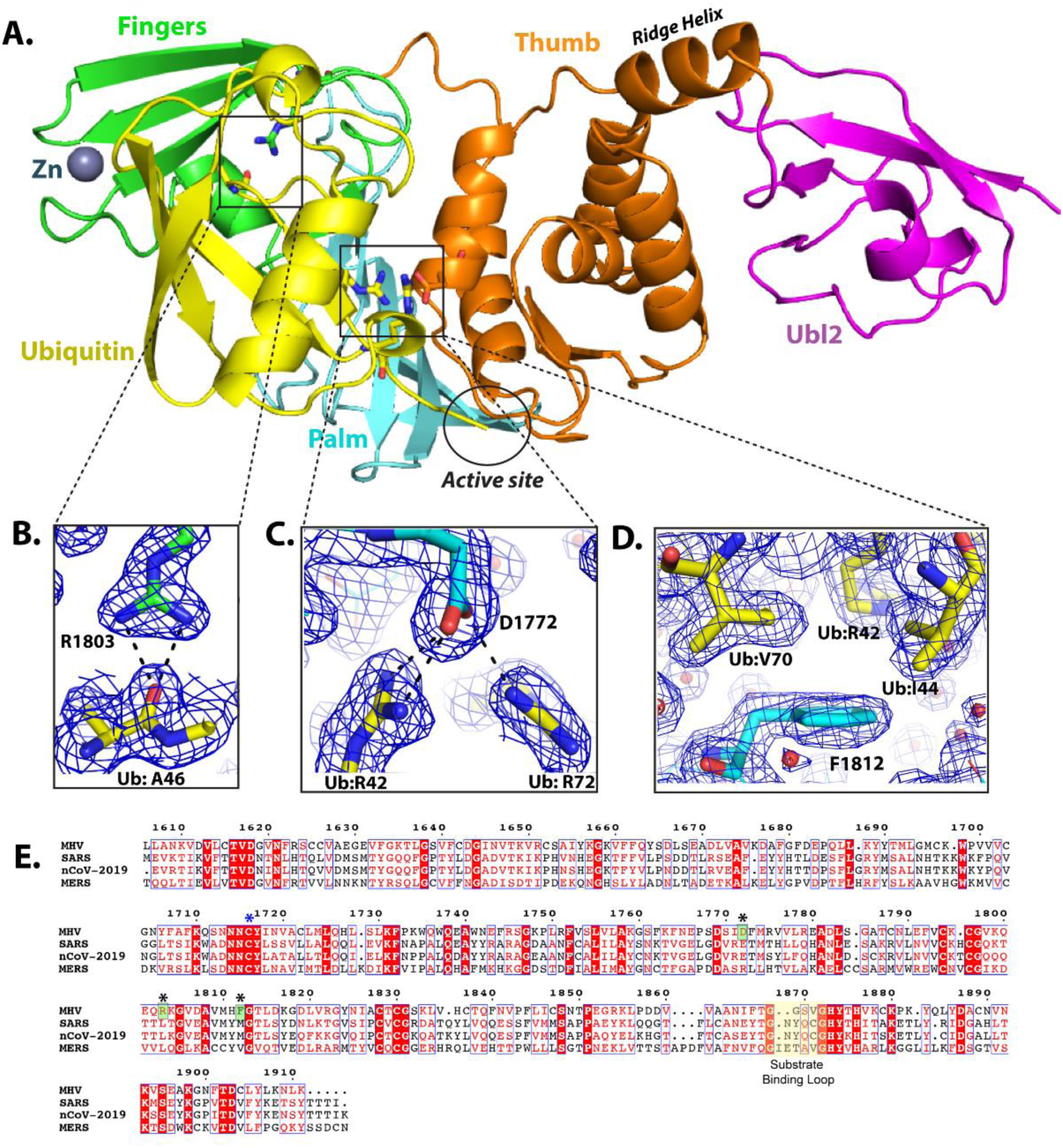
X-ray structure of the MHV PLP2-Ubiquitin complex and residues involved in ubiquitin-binding. (A) Overall structure of MHV PLP2-C1716S-Ub complex. Domains are color coded: Ub in yellow, Ubl2 domain of PLP2 in purple, thumb domain in orange, palm domain in cyan, and fingers domain in green. Residue D1772 of PLP2 is located outside of the active site which is circled in black. (B) Hydrogen bond interactions between MHV PLP2 R1803 and the backbone of A46 of ubiquitin. (C) Binding interactions between MHV PLP2 D1772 and two arginine residues (R42 and R72) of ubiquitin. (D) Hydrophobic interactions between F1812 of MHV PLP2 and ubiquitin residues I44 and V70. The 2Fo-Fc maps (blue) surrounding the residues are contoured at 1s in each panel. The PDB coordinates for the MHV PLP2-C1716S-Ub complex have been deposited under PDB Code 5WFI. (E) Sequence alignment (MultAlign) of coronavirus papain-like protease/deubiquitinating domains from MHV (1606-1911 aa, accession #AAX23975), SARS (1541-1854 aa, accession #ACZ72209), 2019-nCoV (1564-1878, accession #QHO60603) and MERS (1480-1803 aa, accession # AHY21467). Amino acids are colored by similarity using the RISER coloring scheme. Numbering shown is based on MHV sequence. Amino acids mutated in this study are indicated with a black asterisk, the catalytic cysteine is indicated by a blue asterisk, and those amino acids that bind ubiquitin and were mutated in this study are boxed in green. The active site substrate binding loop also involved in binding inhibitors of SARS is shown highlighted in yellow. The sequence alignment was created using ESPript3.

Next, we aligned the primary amino acid sequence of the MHV PLP2 domain with the SARS-CoV and MERS-CoV papain-like protease domains (Fig. 1E). The sequence alignment and X-ray structure of the MHV PLP2–ubiquitin complex were then analyzed in conjunction with the previous structural and mutagenesis studies on SARS- and MERS-CoV to identify candidate residues that could be mutated to render a loss of DUB activity *in vitro*. From this analysis, we identified three residues (R1803, D1772, F1812) in MHV PLP2 that form direct interactions with ubiquitin (Fig. 1B-D). Two of the side chain guanidinium nitrogens of R1803 form direct hydrogen bonds with the backbone carbonyl oxygen of A46 in ubiquitin (Fig. 1B). The two side chain carboxylate oxygens of D1772 in MHV PLP2 interact with ubiquitin by forming direct bonds with each of the guanidinium nitrogens of R42 and with one of the guanidinium nitrogens of R72 (Fig. 1C). Finally, F1812 forms Van der Waals contacts with the side chains of I44 and V70 and the delta-carbon of R42 of ubiquitin (Fig. 1D).

### Biochemical analysis of PLP2 mutants’ activities

To identify a mutant MHV PLP2 enzyme that retains protease activity but exhibits reduced DUB and/or deISGylating activity, we performed site-directed mutagenesis on each of the residues (R1803, D1772, F1812) by changing each to an alanine to disrupt interactions with ubiquitin. Each mutant enzyme was expressed, purified, and tested for its ability to hydrolyze three substrates: z-RLRGG-AMC, Ub-AMC, and ISG15-AMC. The activity of each mutant enzyme toward each substrate relative to the wild-type enzyme is shown in Fig. 2A. All three mutants retained their ability to hydrolyze the peptide substrate but each mutant had altered specificity toward Ub-AMC and ISG15-AMC substrates. Mutation of F1812 resulted in a substantial decrease in hydrolysis of both Ub-AMC and ISG15-AMC (Class I), whereas mutation of R1803 resulted in loss of activity only toward ISG15-AMC (Class II), and mutation of D1772 resulted in loss of activity only toward Ub-AMC (Class III).

**Figure 2.**
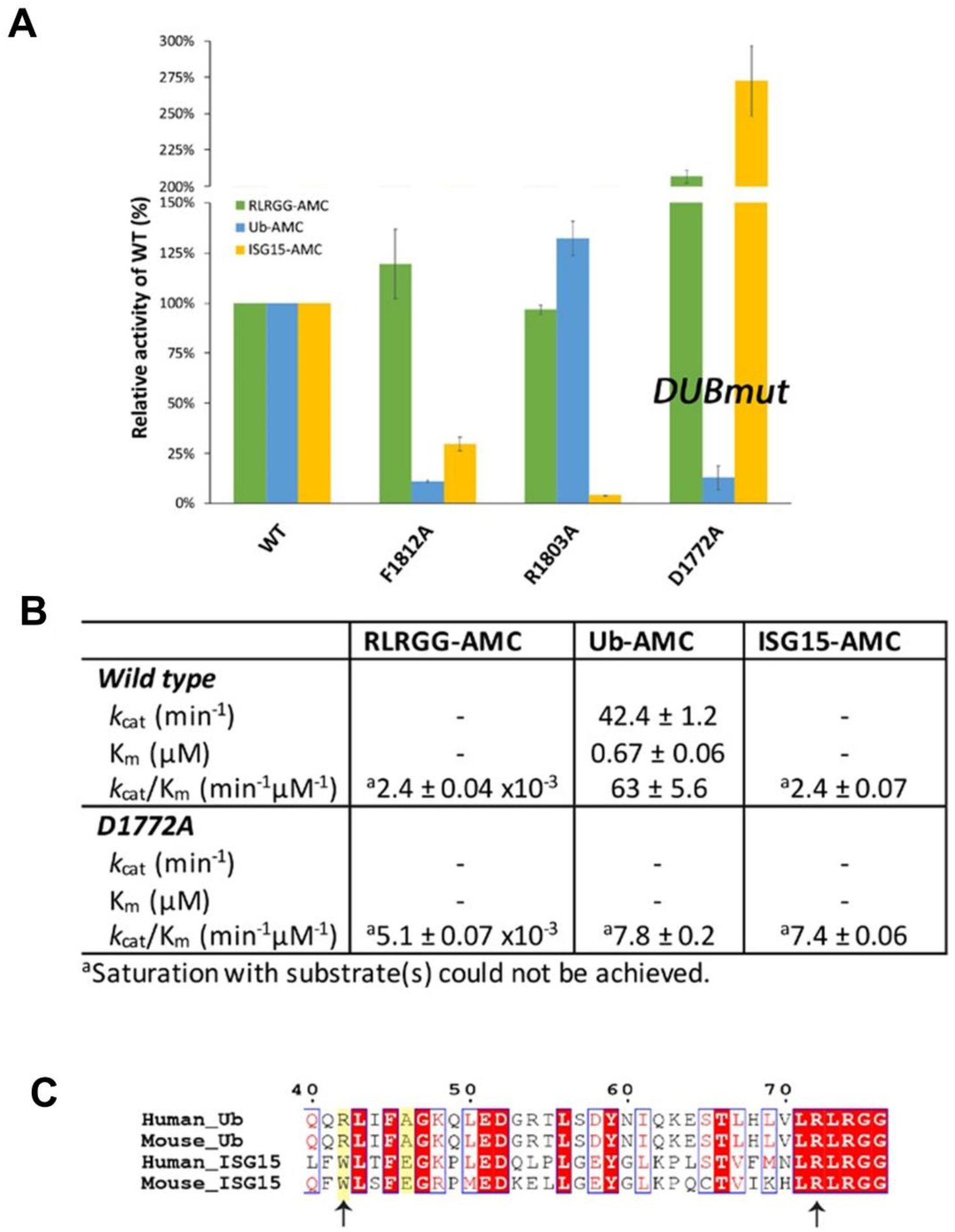
Structure-guided mutagenesis of MHV PLP2 reveals that D1772A disrupts ubiquitin binding and reduces DUB activity. (A) Relative kinetic activities of three mutant MHV PLP2 enzymes toward three substrates: z-RLRGG-AMC (green), Ub-AMC (blue), and ISG15-AMC (yellow) compared to the wild-type enzyme. (B) Steady–state kinetic parameters for wild type and D1772A mutant enzymes. (C) Sequence alignment of Ub and ISG15 from human and mouse generated by Clustal Omega. The two arginine residues of Ub (R42 and R72) that interact with D1772 are indicated by arrows. R72 is conserved between Ub and ISG15, whereas R42 (shaded in yellow) is only present in Ub. Accession numbers: Human_Ub, 1ubq; Mouse_Ub, P62991; Human_ISG15, AAH09507; Mouse_ISG15, AAI09347. The sequence alignment was created using ESPript.

Since one of the primary goals of this study is to understand the contribution of DUB activity to viral replication and pathogenesis, we next focused on quantitating further the effects of the D1772A mutant on the steady-state kinetic parameters of MHV PLP2 toward the three different substrates (Fig. 2B). The RLRGG-AMC peptide substrate is often used as surrogate of the viral polyprotein substrate and the kinetic data in Fig. 2B show that this substrate is still well-recognized and cleaved by the D1772A mutant. In fact, we observed a small rate enhancement in the catalytic efficiency (i.e., *k*_cat_/K_m_) compared to the wild-type enzyme.

The Ub-AMC substrate, on the other hand, is poorly recognized and cleaved by the D1772A mutant compared to the wild-type enzyme. The wild-type enzyme normally interacts strongly with Ub-AMC with a K_m_ value of 0.67 µM. However, mutation of D1772 to an alanine significantly disrupts the interaction with ubiquitin, making it impossible to saturate MHV PLP2 under normal experimental conditions (Fig. 2B). The net result is a significant reduction in the catalytic efficiency (*k*_cat_/K_m_) compared to the wild-type enzyme, which was the goal of these trials.

The kinetic response of MHV PLP2 toward another substrate, ISG15-AMC, was also determined. ISG15 is an important ubiquitin-like modifier that is upregulated and used to ISGylate host proteins during viral infection. A number of viruses, including coronaviruses, engender ISGylation during infection but the function(s) and importance of this activity are not clear (28–30). For MHV, neither the wild-type nor the D1772A mutant PLP2 enzyme can be saturated with ISG15-AMC, suggesting weak binding with this ubiquitin-like modifier (Fig. 2B). Moreover, the R1772A mutation does not disrupt the interaction with ISG15 but in fact enhances it to some degree. A potential explanation for the observed selective disruption of ubiquitin binding stems from our analysis of a primary sequence alignment of ubiquitin and ISG15 and the residues that interact with D1772 (Fig. 2C). The interaction between MHV PLP2 D1772 and the R42 residue in human and mouse ubiquitin is absent in human and mouse ISG15 since this residue is a tryptophan in human and mouse ISG15. Therefore, in line with our observations, D1772A mutation would not be expected to alter ISG15 binding. In contrast to R42, residue R72 is conserved in both ubiquitin and ISG15 and its interaction with MHV PLP2 for ubiquitin is likely weaker than that with ISG15.

### PLP2 D1772A mutant exhibits reduced DUB activity and inhibition of IFN response in cell culture based assays

The *in vitro* biochemical studies presented here support the notion that we are able to use a structure-guided mutagenesis to uncouple the DUB enzymatic activity from MHV PLP2 while preserving the peptide hydrolysis and deISGylating activities of PLP2. Next, we focused on comparing the activity of the mutant enzyme to its wild-type counterpart for the ability to remove Flag-tagged-ubiquitin conjugated to host proteins in cultured cells (Fig. 3A). We found that in cells, wild-type PLP2 exhibits robust DUB activity and removes ubiquitin modifications from multiple cellular proteins. On the other hand, the PLP2-D1772A mutant exhibits reduced DUB activity, similar to that of the previously documented catalytic cysteine to alanine mutant, PLP2-CA (19). To determine if this impaired DUB activity altered the ability of PLP2 to act as an interferon antagonist, we transfected cells with a RIG-I expression plasmid, an interferon-luciferase reporter construct, and either wild-type or mutant PLP2 plasmid and measured luciferase activity at 18 hours post-transfection. In agreement with previous reports (13, 25, 31), we find that wild-type PLP2 acts as an interferon antagonist, reducing reporter activity by 50-80%. In contrast, PLP2-D1772A is unable to significantly reduce interferon activation in this assay despite similar expression levels of the wild-type and mutant versions of the protein (Fig. 3B). We also evaluated the protease activity of the enzymes in cells using two independent *trans*-cleavage assays and found that the wild-type and DUB-mutant enzymes produce similar levels of cleaved products. These results indicate that the D1772A substitution did not alter protease activity (Fig. 3C and D), in agreement with the *in vitro* kinetic results described above (Fig. 2). Together, these studies reveal that aspartic acid residue 1772 of MHV-PLP2 is important for DUB activity and interferon antagonism, but not for protease activity.

**Figure 3.**
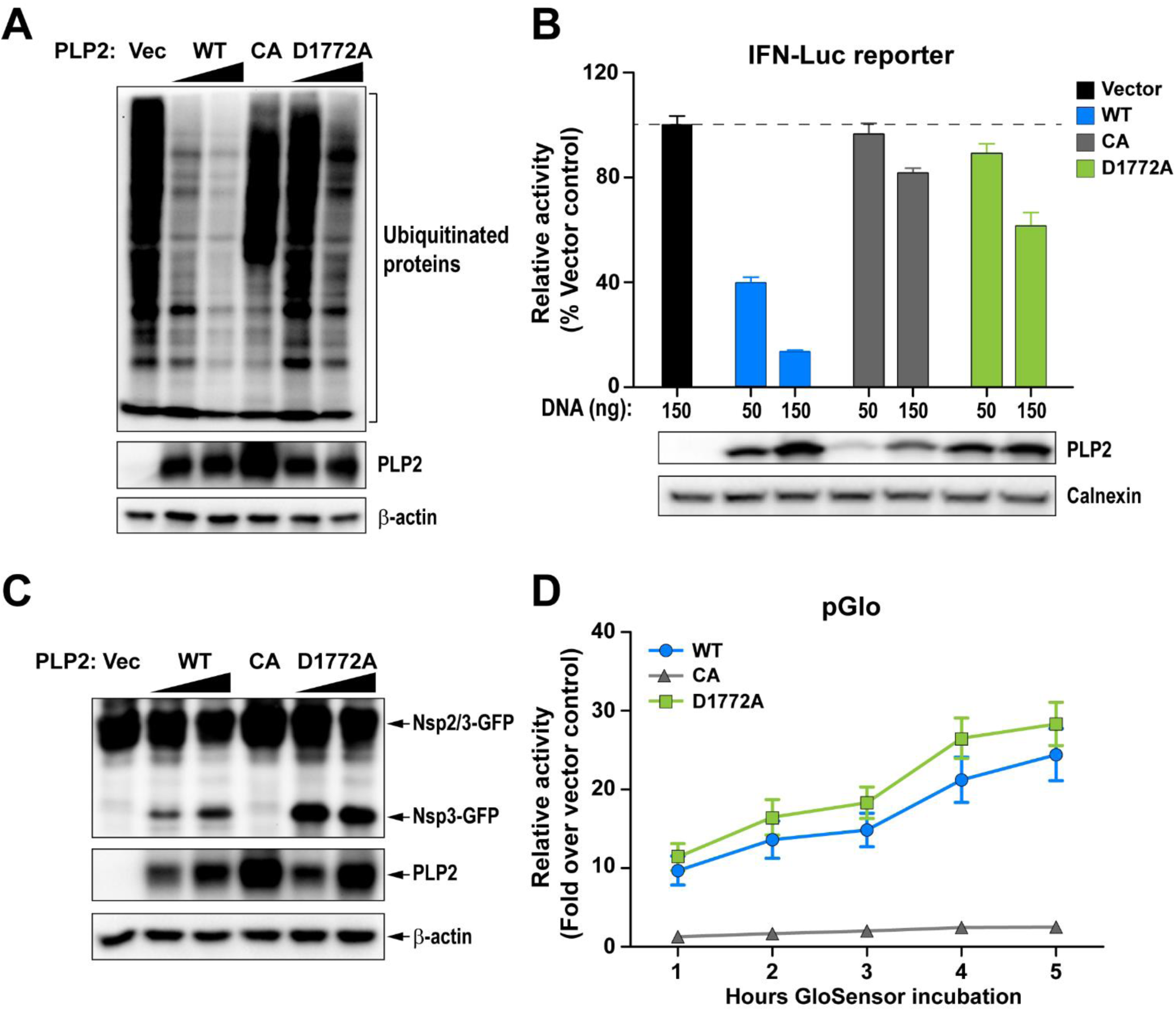
D1772A substitution in the coronavirus papain-like protease Ub-binding site reduces DUB activity and interferon antagonism without reducing protease activity. (A) Western blot assessing the DUB activity of PLP2. (B) IFN antagonism of PLP2 was determined using an IFN-luciferase reporter stimulated by N-RIG-I expression. The reporter activity of vector control was set to 100% (indicated by a dash line). (C and D) Protease activity was evaluated using (C) a trans-cleavage assay that detects the cleaved products by western blot and (D) a pGlo biosensor assay which is activated by PLP2-mediated cleavage of the substrate. Data are representative of at least two independent experiments. Data in (B) and (D) are presented as means ± SD.

### Recombinant MHV harboring PLP2-D1772A activates an earlier IFN response in bone marrow-derived macrophages

Since the D1772A substitution did not impact protease activity, we reasoned that we should be able to generate recombinant virus containing this substitution, thereby allowing us to determine if the mutation has any effect on viral replication kinetics and interferon antagonism in the context of the live virus. We engineered the mutant virus *via* reverse genetics (32), performed full genome sequencing to verify the genotype (2 nucleotide changes at positions 5525 and 5526, resulting in D1772A substitution in the replicase polyprotein), and designated the virus as DUBmut. Upon evaluating virus replication of the DUBmut virus by performing a growth kinetics experiment in parallel with wild-type virus, we found that the DUBmut virus replicates with essentially identical kinetics as the wild-type in a murine astrocytoma cell line (DBT cells) (Fig. 4A). These results are consistent with previous studies of coronavirus interferon antagonists, which showed in many cell lines that viral-mediated interferon antagonism is not essential for virus replication (5, 6). Regarding the other ubiquitin-interacting residues identified in the structural analysis, we attempted to rescue virus with substitutions at the F1812 position, but were unable to recover viable virus. These results indicate that F1812 may play a critical role within the polyprotein during virus replication. We were able to recover virus containing the R1803A substitution, but found that it had no detectable phenotype, which we documented in our previous study (5). Here, we focus our efforts on evaluating replication and pathogenesis of the recovered DUBmut virus.

**Figure 4.**
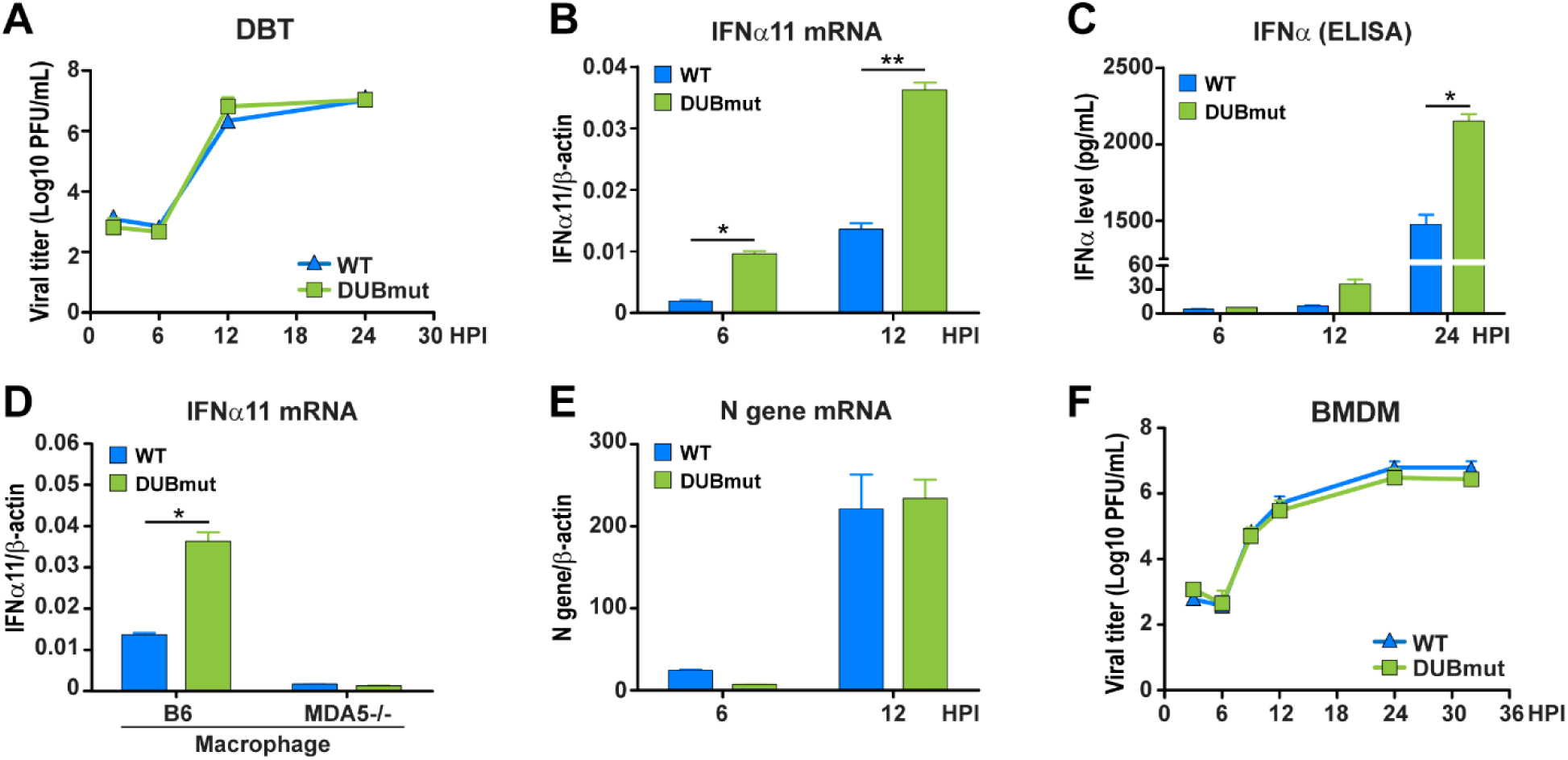
Evaluating the replication kinetics of, and level of interferon activation by, WT MHV and DUBmut in cell culture. (A) Replication kinetics of WT and DUBmut virus in DBT cells. (B) IFNα11 mRNA levels in WT- and DUBmut-infected BMDMs were assessed at indicated time points by qRT-PCR. (C) IFNα protein levels in the supernatants of infected BMDMs were evaluated at the times indicated. (D) Comparison of IFNα11 levels in B6 versus MDA5-/- BMDMs at 12 hours post-infection (HPI). (E) Assessing levels of viral nucleocapsid (N) mRNA by qRT-PCR. (F) Replication kinetics of WT and DUBmut virus in BMDM cells. Data are representative of at least two independent experiments and are presented as means ± SD. Data in (B) and (C) were statistically analyzed using unpaired t-tests. *, p < 0.05; **, p < 0.01.

To determine if the impaired DUB activity of the DUBmut virus had an effect on interferon antagonism, we infected primary bone marrow-derived macrophages (BMDMs) and evaluated viral replication kinetics and levels of interferon mRNA and protein. We observed significant activation of interferon alpha (IFNα) mRNA expression (Fig. 4A) that is coupled with release of IFNα protein into the supernatant, as detected by ELISA (Fig. 4B). We show that this activation of IFNα is dependent on expression of pattern recognition receptor MDA5 (Fig. 4D), in agreement with previous reports (5, 6, 33). To our surprise, we found that replication of DUBmut is not impaired relative to the wild-type virus in BMDMs, as measured by level of nucleocapsid RNA (Fig. 4E) and evaluation of infectious virus particle production over time in the kinetics assay (Fig. 4F). These results demonstrate that an elevated interferon response is generated during replication of the DUBmut virus, but that this interferon profile is not associated with reduced production of infectious particles in either DBT cells or BMDMs.

### Disrupting DUB activity mildly attenuates coronavirus pathogenesis in mice

To complement our *in vitro* studies, we next sought to determine whether loss of DUB activity and the observed activation of interferon during the DUBmut virus infection in macrophages is associated with an attenuated pathogenesis in mice. To test this, we first inoculated mice using the intracranial infection model to evaluate lethality among WT and DUBmut-infected mice, and found similar weight loss and lethality (data not shown). To investigate the pathogenesis in the liver, we inoculated four- (young) or six-week-old (adult) mice intra-peritoneally with a low dose (6×10^3^ plaque-forming units [pfu]) or a high dose (6×10^4^ pfu) of the designated virus, respectively, and measured viral titer in the liver and spleen at the indicated days post-infection (dpi). We observed similar levels of infectious particles in young mice inoculated with a high dose of virus and adult mice with a low dose at 3 and 5 dpi (data not shown), but detected reduced viral titers in the spleens of young mice infected with a low dose of DUBmut virus at 3 dpi (Fig. 5A) and adult mice inoculated with a high dose at 5 dpi (Fig. 5B). Similar pathology in liver sections was observed at 3 and 5 dpi (Fig. 5C). These results indicate that the DUBmut virus is mildly attenuated compared to the wild-type virus, suggesting that the DUB activity does play a role in MHV pathogenesis and PLP2 is a virulence factor.

**Figure 5.**
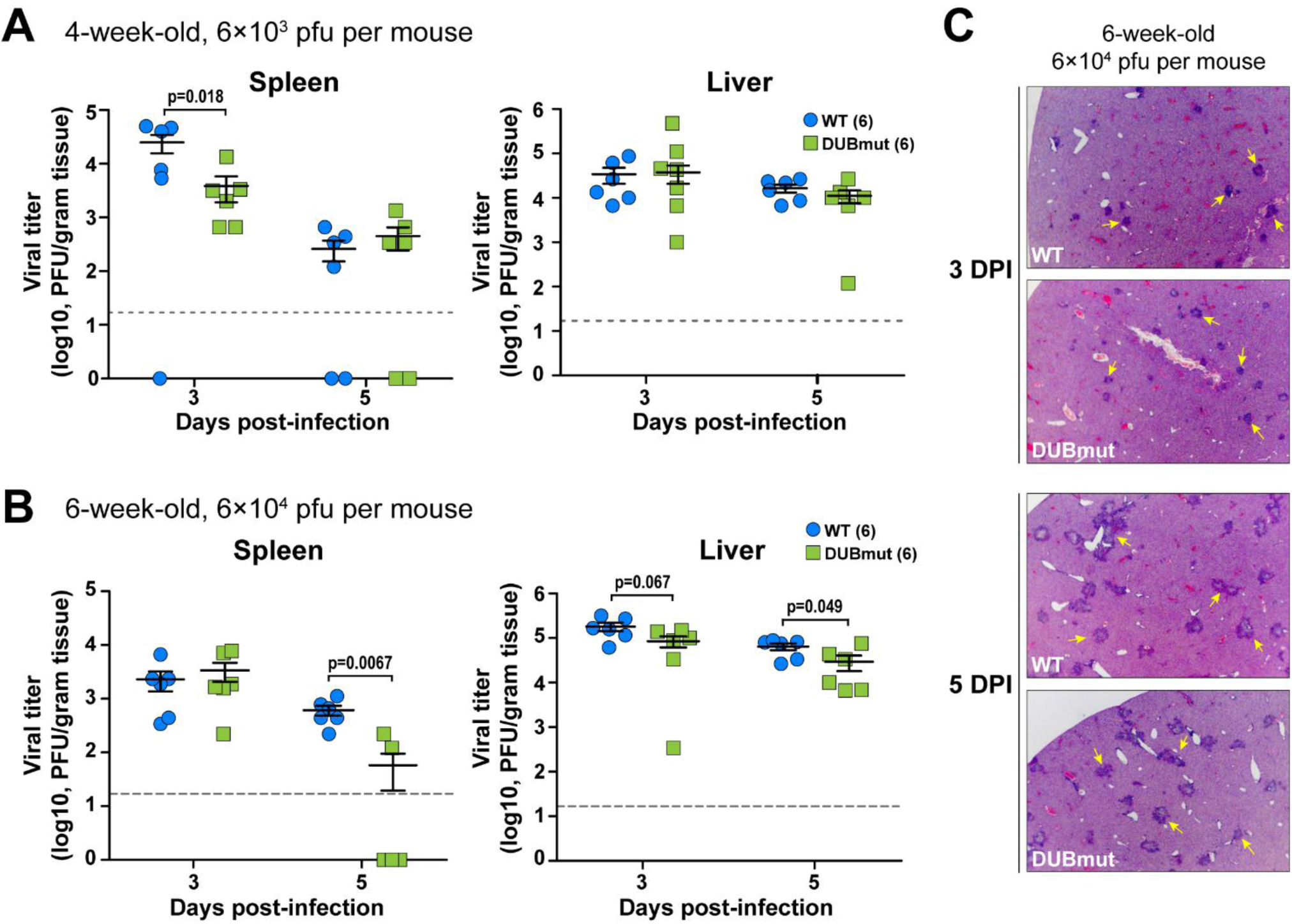
Evaluating replication and pathogenesis of MHV-DUBmut in mice. Four- or six-week-old (B) mice were infected with indicated doses of MHV. Viral titer in livers and spleens isolated from WT- or DUBmut virus-infected mice was determined by plaque assay. The number of mice in each group is shown in parentheses. Data were statistically analyzed using unpaired t-tests and are presented as means ± SEM. (C) H&E staining of liver sections from infected mice at 3 and 5 days post-infection (DPI). Representative MHV-associated liver lesions are indicated by arrows.

## Discussion

In the present study, we aimed to investigate the roles of PLP2 DUB activity during coronavirus infection. Through structure-guided mutagenesis, we identified residues of MHV PLP2 that mediates its interaction with ubiquitin. By mutating the residue Asp-1772 to Ala (D1772A), we found that the DUB activity of PLP2 was greatly reduced and a recombinant MHV carrying this mutation (DUBmut) activated an earlier IFN response in macrophages. Although we only observed a subtle attenuation of the DUBmut virus in the tested animal models, we demonstrated that PLP2 DUB activity does play a role in suppressing the host immune response and is a virulence trait, strengthening our earlier discoveries on SARS-CoV PLpro (4). We further investigated the differences in the interferon response generated in response to WT versus DUBmut virus in our companion study (Volk et al., submitted), which further supports a role for DUB activity in modulating the interferon response in macrophages.

The structure-guided approach used to generate the DUBmut virus allowed for characterization of three different classes of mutant enzymes: Class I, deficient in both DUB and deISGylating activity; Class II, deficient in deISGylating activity only; and Class III, deficient in DUB activity but competent in protease and deISGylating activity. We utilized three unique biochemical substrates, each with a conjugated fluorescent AMC reporter, to evoke the multi-functional activities of PLP2. Activity against the z-RLRGG-AMC peptide substrate represents the polyprotein processing activity of MHV PLP2, while UB-AMC and ISG15-AMC stimulate the deubiquitinating and deISGylating activities of the enzyme, respectively (17, 34). The kinetic data for the D1772A DUBmut hydrolysis of the z-RLRGG-peptide provided in Figure 2B indicate that the polyprotein processing ability of this mutant is likely not affected by the D1772A substitution. The deISGylating ability of the enzyme is also not affected. In contrast, the mutant enzyme’s deubiquitinating activity is significantly reduced relative to wild-type, which is most likely due to a lowered binding affinity for ubiquitin as the enzyme could not be saturated with Ub-AMC as a substrate.

We can also use these structural, mutagenesis and kinetic characterization studies on MHV PLP2 to guide future structure-based design studies of emerging coronaviruses, such as the newly emerged novel coronavirus 2019 (nCoV-2019) (35–38). By aligning the X-ray structure of the PLpro domains of SARS-CoV bound to ubiquitin (17) with the sequence of nCoV-2019 PLpro (Fig. 6A) and comparing the differences in their residues with those of MHV PLP2 (Fig. 1E), we observe significant differences in the potential interactions between nCoV-2019 and ubiquitin. For example, we found that the hydrogen bonds between MHV PLP2 R1803 and the carbonyl oxygen of UbA46 (Fig. 1B) are lost in nCoV-2019 and SARS-CoV (Fig. 6B). In contrast, the hydrogen bonds between MHV PLP2 D1772 and Ub residues R42 and R72 (Fig. 1C) are likely preserved with the substitution of E1772 in nCoV-2019 and SARS-CoV PLpro (Fig. 6C). The hydrophobic interactions between Ub residues I44 and V70 and F1812 of MHV PLP2 (Fig. 1D) also appear to be preserved with M1812 of nCoV-2019 and SARS-CoV PLpro (Fig. 6D). The X-ray structures of MHV PLP2 and SARS-CoV PLpro, together with our structural model of nCoV-2019 PLpro, will provide testable hypotheses for the design and mutagenesis of the ubiquitin-binding domain (17) and diubiquitin-binding domains (20) of the recently emerged coronavirus nCoV-2019 PLpro.

**Figure 6.**
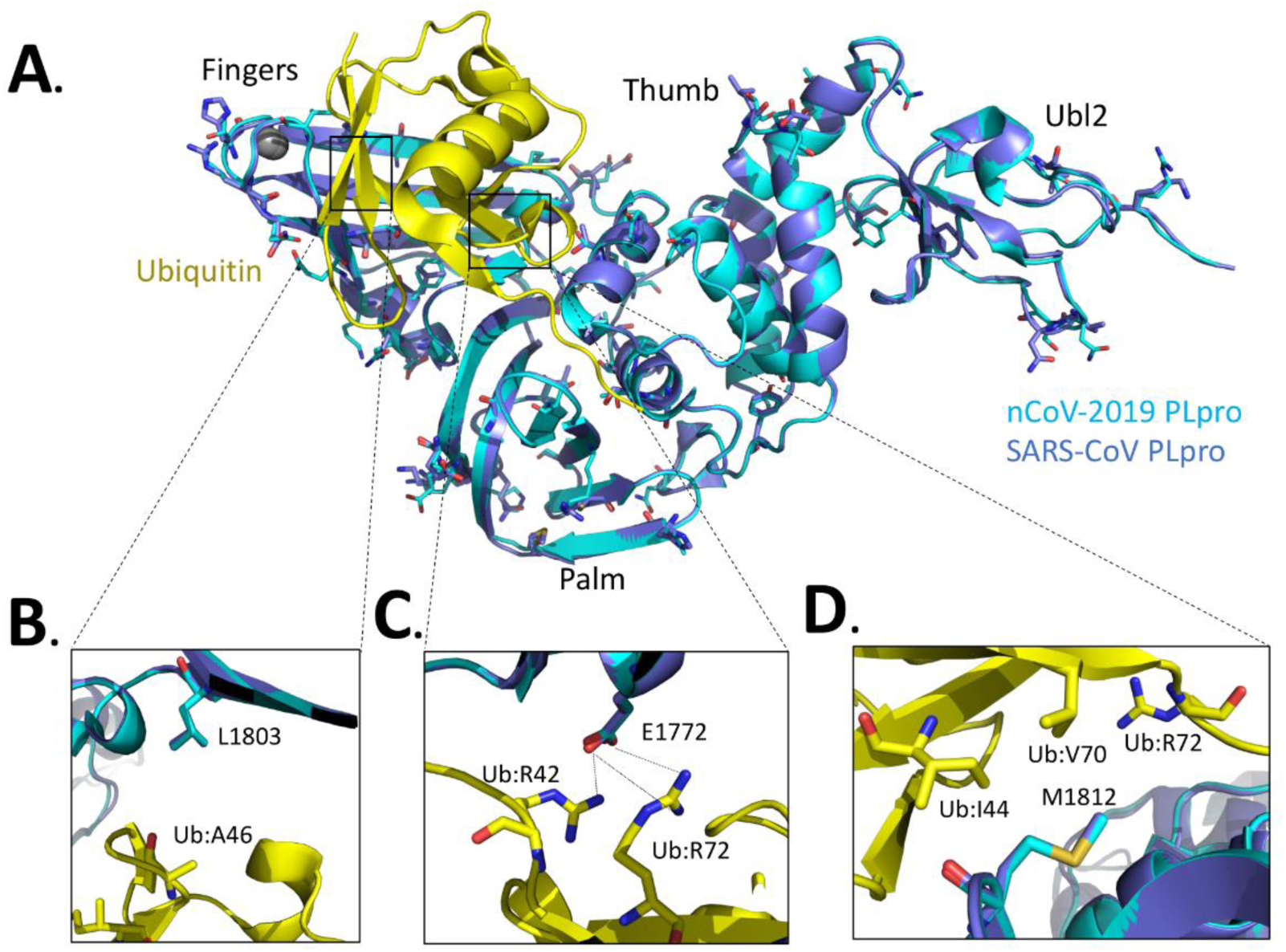
Alignment of the nCoV-2019 PLpro domain with the X-ray structure of the closely related SARS-CoV PLpro domain in complex with ubiquitin. (A) X-ray structure of SARS-CoV PLpro-ubiquitin-aldehyde complex (blue) (PDB: 4MM3) with each of its domains labeled as finger, palm, thumb, and Ubl2. Ubiquitin-aldehyde is colored yellow. The nCoV-2019 PLpro structure (cyan) was modeled by first mutating the residues of SARS PLpro in the X-ray structure to those of nCoV-2019 PLpro based upon the sequence alignment in Figure 1E. The nCoV-2019 PLpro-Ubiquitin-aldehyde complex was then refined using the structure-factor amplitudes and initial phases of the SARS PLP-Ubiquitin aldehyde complex (PDB: 4MM3). The residues that are different between SARS-CoV PLpro and nCoV-2019 PLpro are highlighted as sticks. (B) Potential interactions between L1803 of SARS-CoV PLpro and nCoV-2019 PLpro with residue A46 of ubiquitin. (C) Predicted interaction between E1772 of SARS-CoV PLpro and nCoV-2019 PLpro and residues R42 and R72 of ubiquitin. (D) Potential interactions between residues I44, V70, and R42 of ubiquitin with residues M1812 of SARS-CoV PLpro and nCoV-2019 PLpro.

We were able to reproduce the enzymatic profile of the purified PLP2-D1772A mutant protein when we expressed it in cell culture (Fig. 3). Therefore, our finding that the DUBmut virus containing the PLP2-D1772A substitution activates an elevated antiviral response in macrophages compared to the wild-type virus, but that this antiviral state results in only mild attenuation of disease in mice relative to WT infection, was unexpected. Previous studies demonstrated that ubiquitin has important roles in both the activation and the attenuation of innate antiviral pathways (39); therefore, we anticipated a more remarkable phenotype for a DUB-mutant virus. We can imagine several possible explanations for our findings. First, it is possible that viral DUB activity has a relatively minor role in shaping pathogenesis in this system. In fact, our recent studies using SARS-CoV and SARS-related CoVs found that the papain-like protease domain/DUB is a virulence trait that varies among members of the SARS-coronavirus species (4). In that study, we found that replacing the SARS PLP2/DUB domain with a SARS-related PLP2/DUB domain reduced the ability of that virus to antagonize the innate immune response. Together, this previous report in conjunction with the current study support the concept that different PLP2/DUB domains may have distinct effects on antagonism of the innate immune response depending on the virus and the host cell type. Another possibility is that the DUB-mutant virus we generated may not have been sufficiently debilitated in its DUB activity to result in altered pathogenesis. We found that it was difficult to recover viable DUB-mutant viruses; indeed, this D1772A mutant was the sole viable DUB-mutant representative of our many attempts. Because an elevated interferon response was elicited from the DUBmut-infected cells, this mutant virus fulfilled our criteria demonstrating the inactivation of an interferon antagonist. However, we speculate that if we are able to recover mutants that exhibit a range of DUB activity, we may be able to more fully assess the role of DUB activity as a contributor to coronaviral pathogenesis. Despite these caveats, the MHV DUB-mutant generated in this study did exhibit a reproducible phenotype of eliciting an elevated interferon response in infected macrophages that was associated with mild attenuation of pathogenesis with reduced titers in the livers and spleens of infected mice. Overall, we conclude that DUB activity is indeed a virulence trait, and contributes to the ability of MHV to modulate the host innate immune response to infection. Further studies are needed to identify the targets of viral DUB activity and the detailed role of DUB activity in delaying the innate immune response to virus replication.

## Materials and Methods

### Ethics Statement

The mouse experiment in this study was carried out in accordance with the recommendations in the Guide for the Care and Use of Laboratory Animals of the National Institutes of Health. The experimental protocol was reviewed and approved by the Institutional Animal Care and Use Committee (IACUC) at Loyola University Chicago (IACUC#: 2016-029). C57BL/6 female mice were purchased from The Jackson Laboratory and maintained in the Comparative Medicine Facility of Loyola University Chicago. Mice were consistently monitored for signs of distress over the course of the experiments to be removed from the experiment and euthanized using carbon dioxide inhalation to prevent unnecessary suffering.

### Cells

Human embryonic kidney (HEK) 293T cells were purchased the from American Type Culture Collection (ATCC, # CRL-11268) and maintained in DMEM (#10-017-CV, Corning) containing 10% fetal calf serum (FCS) and supplemented with 1% nonessential amino acids, 1% HEPES, 2% L-glutamine, 1% sodium pyruvate, and 1% penicillin/streptomycin. DBT cells were cultured in MEM (#61100-061, ThermoFisher) supplemented with 5% FCS, 2% L-glutamine, and 10% Tryptose Phosphate Broth (TPB). The 17Cl-1 cell line was maintained in DMEM containing 5% FCS. Baby hamster kidney cells expressing the MHV receptor (BHK-R) were kindly provided by Mark Denison (Vanderbilt University Medical Center) and maintained in DMEM supplemented with 10% FCS, 2% L-glutamine, and 0.8 µg/mL G418. Bone marrow-derived macrophages (BMDMs) were prepared and cultured as described previously (5).

### Plasmids and mutagenesis

The sequence of the PLP2 domain (amino acids 1525-1911 of MHV pp1ab) in frame with a V5 epitope tag was codon-optimized, synthesized by Genscript (Piscataway, NJ) (sequence available upon request), and cloned into pCAGGS vector. For mutagenesis, an overlapping PCR strategy was used with synthetic primers (sequences available upon request). The introduced mutations were verified by sequencing. The RIG-I and nsp2/3-GFP expression plasmid was kindly provided Ralph Baric (University of North Carolina). The IFNβ-Luc reporter plasmid was a gift of John Hiscott (Jewish General Hospital, Montreal, Canada). The Flag-Ub plasmid was kindly provided by Adriano Marchese (University of Wisconsin-Milwaukee).

### MHV PLP2 wild-type and mutant purification, kinetics, and X-ray structure

The wild-type, C1716S, D1772A, R1803A, and F1812A mutant enzymes were expressed and purified similar to our previously published methods except that the MHV PLP2 construct used here (amino acids N1609 to N1909) was absent the DPUP domain (16). Crystallization and X-ray structure determination details will be published elsewhere. Steady-state kinetic studies on the wild-type, D1772A, R1803A, and F1812A mutant enzymes with substrates Z-RLRGG-AMC (where Z is a carboxybenzyl protecting group), Ubiquitin-AMC, and ISG15-AMC were performed as described previously (16). Structure figures were generated with the software program UCSF Chimera (40).

### Protease and deubiquitinating activity assays

To determine the protease activity of PLP2, HEK293T cells grown to 70% confluency in 24-well plates (Corning) were transfected using *Trans*IT-LT1 (MIR2300, Mirus) according to the manufacturer’s protocol. For the protease activity assay, HEK293T cells were transfected with 25 ng nsp2/3-GFP plasmid and 200 ng pCAGGS-PLP2-V5 expression plasmids (wild-type and mutant). To determine deubiquitinating activity of the proteins, cells were transfected with 200 ng Flag-Ub plasmid and pCAGGS-PLP2-V5 expression plasmids (wild-type and mutant). Cells were lysed 24 h post-transfection with 100 µL of lysis buffer (comprising 20 mM Tris [pH 7.5], 150 mM NaCl, 1mM EGTA, 1mM EDTA, 1% Triton X-100, 2.5 mM sodium pyrophosphate, 1mM β-glycerophosphate, 1mM sodium orthovanadate, 1 µg/mL leupeptin, and 1mM phenylmethylsulfonyl fluoride). Whole cell lysates were separated by SDS-PAGE and transferred to PVDF membrane in transfer buffer (0.025M Tris, 0.192M glycine, 20% methanol) for 1 hour at 60 Volts at 4°C. Following this, the membrane was blocked using 5% dried skim milk in TBST buffer (0.9% NaCl, 10mM Tris-HCl, pH7.5, 0.1% Tween 20) overnight at 4°C. The membrane was incubated with either polyclonal rabbit anti-GFP antibody (A11122, Life Technologies) for the protease assay, or mouse anti-flag (F3165, Sigma) for the DUB assay. The membrane was then washed three times for 15 minutes in TBST buffer followed by incubation with either secondary donkey anti-rabbit-HRP antibody (711-035-152, Jackson ImmunoResearch) or goat anti-mouse-HRP antibody (1010-05, SouthernBiotech). Then the membrane was washed three more times for 15 minutes in TBST buffer. Detection was performed using Western Lighting Chemiluminescence Reagent Plus (PerkinElmer) and visualized using a FluoroChemE Imager (Protein Simple). The expression of PLP2, β-actin, and calnexin were probed with mouse anti-V5 antibody (R960, ThermoFisher), mouse anti–β-actin (A00702, Genscript), or mouse anti-calnexin antibody (2433S, Cell Signaling), respectively.

### Biosensor live cell assay

The protease activity of PLP2 was also assessed using a biosensor live cell assay as described previously (14). Briefly, HEK293T cells in a 96 black-wall plate were transfected with 37.5 ng pGlo-RLKGG construct and 50 ng PLP2 expression plasmids. GloSensor (E1290, Promega) reagent diluted in DMEM+10% FCS was added at 18 hpi. Plates were read using a luminometer (Veritas) every hour over a course of 5 hours.

### Generating DUB-mutant MHV

We used a previously described reverse genetics system of MHV-A59 (32) to generate the DUB-mutant virus. Briefly, the nucleotides coding for the Asp-1772 of the PLP2 domain were mutated *via* site-directed mutagenesis. Viral genomic RNA from *in vitro* transcription of ligated cDNA fragments using a mMESSAGE mMACHINE T7 Transcription Kit (AM1344, Thermal Fisher) was electroporated into BHK-R cells. Cell supernatants were collected as viral stock following observation of cytopathic effects. Rescued virus was plaque-purified, propagated on BHK-R cells, and titrated on 17Cl-1 cells. The stock virus was subjected to full-genome sequencing and the sequences were aligned to the parental strain, with the intended engineered mutation detected and no additional mutations detected (Kansas State University Diagnostic Laboratory).

### Growth kinetics

DBT cells or BMDMs were infected with wild-type icMHV-A59 or DUB-mutant virus at a multiplicity of infection (MOI) of 1 in serum-free medium. After a one-hour incubation, the inoculum was replaced with fresh, complete medium. Cell culture supernatants were collected at indicated time points and titrated by plaque assay on 17Cl-1 cells. Titers were obtained from three independent assays for each sample. Graphs of virus kinetics were generated using Prism software (GraphPad Software).

### Quantification of IFNα production by RT-qPCR and ELISA

BMDMs in a 12-well plate were mock-infected or infected with MHV at a MOI of 1. At indicated time points, monolayer cells were collected for RNA extraction and cell culture supernatants were harvested for ELISA analysis. To determine IFN-α11, β-actin, or MHV-A59 N gene mRNA levels, total RNA was extracted from collected cells using a RNeasy Mini Kit (74104, Qiagen). The first strand cDNA was synthesized from an equal amount of RNA using Rt2 HT First Strand Kit (330401, Qiagen). qPCR was then performed with specific primers for mouse IFN-α11 (PPM03050B-200, Qiagen), mouse β-actin (PPM02945B-200, Qiagen), or MHV-A59 N gene using RT2 SYBR Green qPCR Mastermix (330502, Qiagen) in the Bio-Rad CFX96 system. The thermocycler was set as follows: one step at 95°C (10 min), 40 cycles of 95°C (15 s), 60°C (1 min) and plate read, one step at 95 °C (10 s), and a melt curve from 65°C to 95°C at increments of 0.5°C/0.05s. Samples were evaluated in triplicate and data are representative of three independent experiments. The levels of mRNA were reported relative to β-actin mRNA and expressed as 2^−*ΔCT*^ [*ΔC*_*T*_ *= C*_*T(gene of interest)*_ − *C*_*T(β-actin)*_]. The secreted amount of IFN-α in culture supernatants was assayed using a mouse IFN-α ELISA kit (BMS6027, eBioscience) according to the manufacturer’s instructions.

### Mouse experiments

Evaluation of MHV pathogenesis in laboratory mouse was previously described (5, 41). Briefly, for intracranial infections, six-week-old C57BL/6 female mice were inoculated with 600 pfu of virus in 20 μL PBS. Infected mice were monitored for body weight daily and euthanized when weight loss surpassed 25%. Statistical analyses of survival rate were performed using the log-rank test. For intraperitoneal infection, six- or four week-old mice were injected with 6×10^3^ or 6×10^4^ pfu of virus in 100 μL PBS. Organs were collected at indicated time points and evaluated for viral burden. Liver pathology was evaluated using H&E staining by the Tissue Processing Core Facility at Loyola University Chicago.

## Acknowledgments

This work was supported by the National Institutes of Health (NIH) grant R01 AI085089 (to SCB and ADM). MH and RCM were supported by NIH T32 Training Grant for Experimental Immunology (#AI007508) and RCM was supported by the Arthur J. Schmitt Dissertation Fellowship in Leadership and Service (Arthur J. Schmitt Foundation). MEC was supported by an NIH/NIGMS T32 Training Grant for Structural Biology and Biophysics (#GM132024). Crystallization and DNA sequencing were partially supported by the Purdue Center for Cancer Research Macromolecular Crystallography and DNA Sequencing Shared Resources, which are supported by NIH grant P30 CA023168.

## Author Contributions

Conceptualization: XD, ADM and SCB. Investigation: XD, YC, AMM, MH, KRK, RCM, MH, AO, MEC. Formal Analysis: XD, YC, AMM, MH, KRK, RCM, AO, ADM and SCB. Writing – Original Draft Preparation: XD, YC, AMM, ADM and SCB. Writing – Review & Editing: with comments from XD, YC, AMM, MH, KRK, RCM, MH, AO, ADM and SCB. Visualization: XD, YC, KRK, AO, MEC, ADM and SCB. Funding Acquisition and Supervision: ADM and SCB.

